# Patterned 3D printed hydrogel as a novel soilless substrate for plant cultivation

**DOI:** 10.1101/2025.08.06.668759

**Authors:** Ali Mohammed, Maddalena Salvalaio, Yumeng Li, Connor Myant, Giovanni Sena

## Abstract

Plant roots need water, micronutrients, and oxygen to maintain cellular metabolism and tissue growth, yet traditional hydroponic systems often lack sufficient oxygen delivery. While 3D printing artificial substrates has been explored to mimic the physical structure of soil, it remains unclear which design parameters are critical for supporting full plant development. Here, we present a synthetic, soilless substrate based on 3D-printed hydrogels incorporating triply periodic minimal surface (TPMS) patterns to create internal air-filled channels. These channels are open to the atmosphere, enabling passive gas exchange throughout the substrate.

We tested five TPMS geometries (Lidinoid, Split-P, Schwarz-D, Schwarz-P, and Schoen), each with a near-identical hydrogel volume but with different surface-to-volume ratios. *Arabidopsis thaliana* seeds germinated directly on the substrates and were monitored for vegetative and reproductive growth over five weeks. Among the designs, the Lidinoid substrate led to the highest number and surface area of leaves, exceeding the aerated hydroponic control in leaf area over most of the time course; it also produced the earliest and most complete flowering, which neither control achieved in the same amount of time.

Our results indicate that the surface-to-volume ratio is a key parameter influencing substrate performance; we hypothesise that this reflects its effect on oxygen availability at the root interface, which we did not measure directly. Plants grown on substrates with higher surface areas transitioned to flowering more reliably and rapidly, with flowering efficiency showing a strong positive correlation with surface area. These findings suggest that interconnected air-channel architectures can support full plant development without active aeration, potentially by overcoming the oxygen limitations of traditional hydroponic systems. This work supports the use of additive manufacturing as a powerful tool for engineering soil analogues tailored for indoor agriculture. By combining passive aeration with hydration and nutrient delivery, patterned hydrogels offer a promising, scalable solution for sustainable soilless plant cultivation.

## INTRODUCTION

Land plants rely on their root system for water and micronutrient uptake. Root cells depend on oxygen for aerobic cellular respiration, which provides energy in the form of ATP for growth, biomass maintenance, and ion uptake [1,2]. Cellular respiration rates vary with plant species, root tissue [3], and environmental conditions [1], but in all cases access to gaseous oxygen is advantageous; sub-optimal (hypoxia) or undetectable (anoxia) oxygen concentrations diminish aerobic metabolism [4], and prolonged exposure causes severe cellular damage through acidification and accumulation of reactive oxygen and nitrogen species [5,6]. Consequently, oxygen availability in the root zone is crucial for root function and plant productivity, whether in soil or soilless cultivation systems.

Oxygen is poorly soluble in water (Henry’s constant of gaseous vs. aqueous concentration of O₂ at 25 °C and 1 atm is 31.46 [1]) and diffuses approximately 10,000 times faster in air than in water [7]; consequently, air-filled pores represent the primary source of oxygen for roots in soil. Soil aeration depends on diffusion from the atmosphere through open passages [8], and is determined by macroscopic geometry: particle size distribution (soil texture), porosity (empty spaces), pore size distribution, and connectivity of the pore network (coordination number) [1]. Soil can therefore represent an adequate environment for root growth, provided its structure is maintained to preserve aeration.

However, maintaining optimal soil structure over large agricultural areas is challenging and costly. Compaction reduces root growth and plant biomass, due in part to poor soil aeration [9,10]. Various methods have been adopted to counteract this, with mixed results: tillage (e.g. ploughing), air injection, irrigation with water containing air bubbles or hydrogen peroxide, and even the mixing of solid peroxides to soil [1]. Furthermore, conventional soil-based agriculture requires extensive land areas and substantial water for irrigation, which are increasingly scarce and expensive in many regions [11].

These constraints have driven increasing interest in soilless culture systems, which offer potentially greater control over the root environment while reducing land and water requirements. Although liquid (hydroponics and aquaponics solutions) and mist (aeroponics) media can be used to grow small crops [12], the infrastructure and energy consumption required to run and maintain these systems, including pumps and aeration equipment, are often complex and expensive, reducing their commercial and ecological sustainability [13]. Solid soilless substrates such as peat, coir, perlite, and rockwool offer intermediate solutions, providing both structural support and improved aeration compared to liquid culture. However, organic substrates decompose over time and raise environmental concerns, particularly the ecological impact of peat extraction [13,14]. Inorganic substrates have inconsistent pore structures and pose disposal challenges [14]. Crucially, none of these conventional substrates allows precise control over pore geometry, size distribution, or network connectivity; these are parameters that directly influence oxygen diffusion and root-substrate interaction.

Ideally, a soilless substrate should be manufactured to deliver water, nutrients, and oxygen to plant roots passively (i.e., without pumps), while offering control over its internal geometry. The idea of producing a solid substrate that resembles some of the geometrical and mechanical properties of soil, and potentially aeration by diffusion, is not new. A simple approach is to cluster solid fragments randomly, such as in the “transparent soil” made of clear gel [15] or other polymers [16]. More complex structures have been proposed, based on wet granular medium [17] or built with additive manufacturing (3D printing) using hydrogels [18–21] or other materials. In some cases, the 3D-printed structure has been explicitly designed to resemble geometrical traits of real soil [22,23]. Although hydrogels have been 3D-printed using various methods and chemical compositions, not all printable solutions are non-toxic and biocompatible [24].

Overall, these and other results suggest that hydrogel can be adopted as a substrate for plant culture [25] and that a solid substrate can be designed and manufactured to mimic some key properties of soil. It has recently been shown that 3D printed porous substrates made of hydrogel result in increased root growth in some plant species when compared to a non-porous hydrogel substrate [21]. Although this is a significant step forward, a demonstration that the aeration in these designs is sufficient to support sustained and robust plant growth in comparison to traditional hydroponics is, to our knowledge, still missing. Compared to conventional substrates such as peat or coir, 3D-printed hydrogels offer several potential advantages: precise and reproducible control over pore geometry and network connectivity, batch-to-batch consistency, reduced environmental impact (avoiding peat extraction), and optical transparency enabling direct observation of root development. Furthermore, unlike liquid hydroponic media, a patterned hydrogel provides mechanical support and geometrical structure that more closely resembles the heterogeneous environment of real soil.

We present a novel method for designing, optimising, and 3D printing hydrogels with interconnected channel networks that remain in contact with the atmosphere. The mechanical impedance of the hydrogel was chosen to induce the plant roots to grow in the channels without penetrating its volume. We reasoned that this configuration would allow the roots to uptake water and nutrients from the hydrogel medium and, at the same time, avoid hypoxia by being exposed to oxygen at atmospheric concentrations.

We validated the method using *Arabidopsis thaliana* as a model system, showing germination and quantifying leaf size, leaf number, and flowering time across different geometries of the channel network.

## METHODS

### Design

#### TPMS patterns

Among the 3D-printable designs that provide interconnected patterns, we focused on a class of Triply Periodic Minimal Surfaces (TPMS) defined by high surface-area-to-volume ratios (SA/V) and composed of infinite, non-self-intersecting, and periodic surfaces in three principal directions [26,27]. Each TPMS is characterised by a different SA/V. A second important parameter is the percentage of the entire substrate that is composed of hydrogel, referred to as the Hydrogel Fraction (HF). For simplicity, we normalised SA/V to the control case of a cubic substrate with the same dimensions but no channels (referred to as “solid”), defined as SA/V = 1 and HF = 1.

We identified 5 candidate TPMS patterns that, in principle, could be realised across a range of SA/V vs HF (Fig. 1 and 2): Lidinoid, Split-P, Schwarz-D, Schwarz-P and Schoen [27,28]. In practice, spatial resolution limitations in the 3D printing method, together with the structural constraints of preventing the substrate from collapsing under its own weight and sustaining mechanical stress during handling, led us to limit our study to realisations with HF ≈ 0.6 (in the range 0.587–0.608). This value also provides sufficient volume to maintain the internal void structures without becoming solid.

**Fig. 1.**
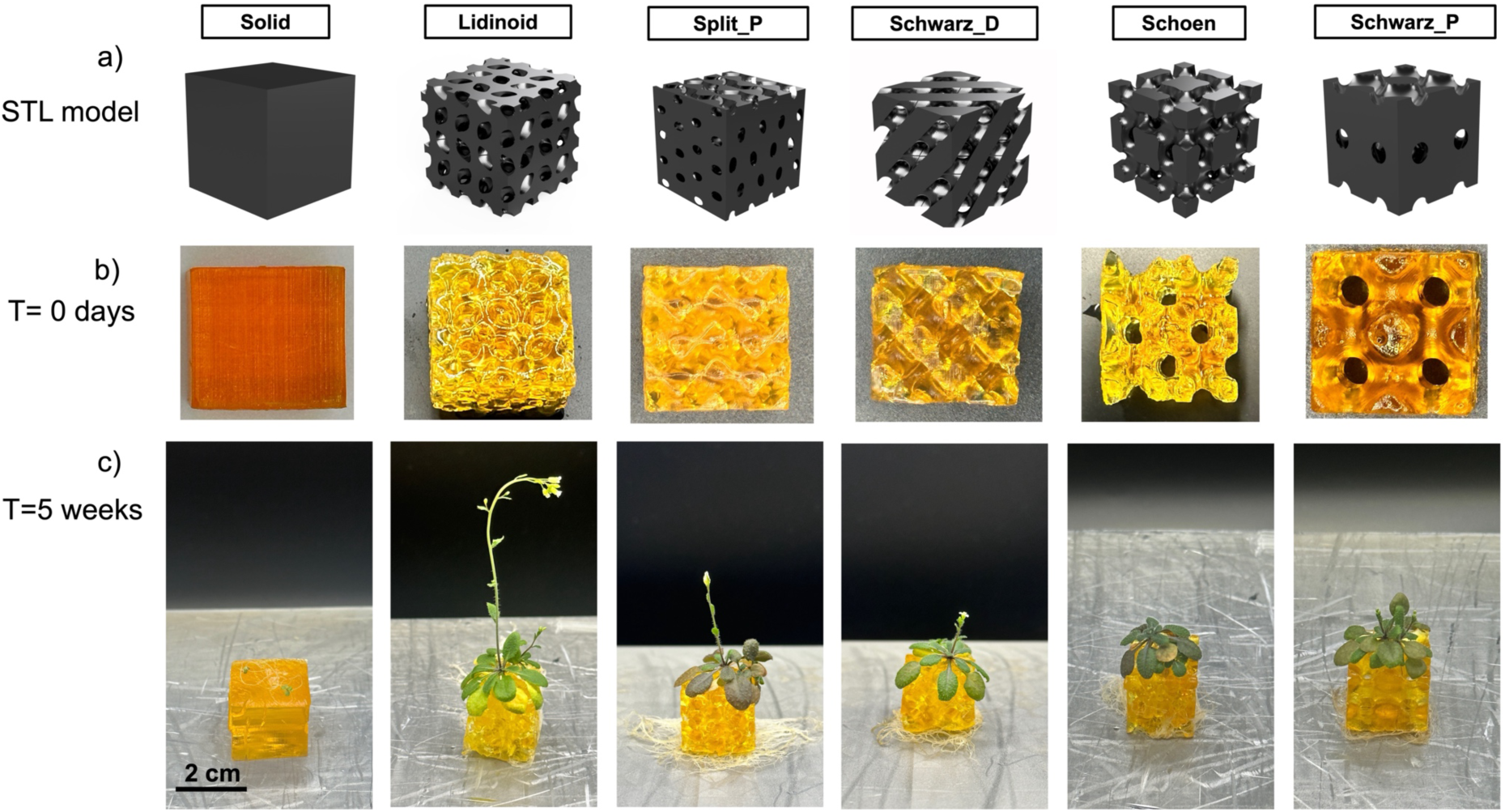
Models and realisations of the 3D-printed hydrogel substrates with each of the tested designs (TPMS patterns and the solid control). **a)** 3D view of the CAD model. **b)** Top view of the printed hydrogel. **c)** Arabidopsis plants germinated and grew for 5 weeks on each design.

**Fig. 2.**
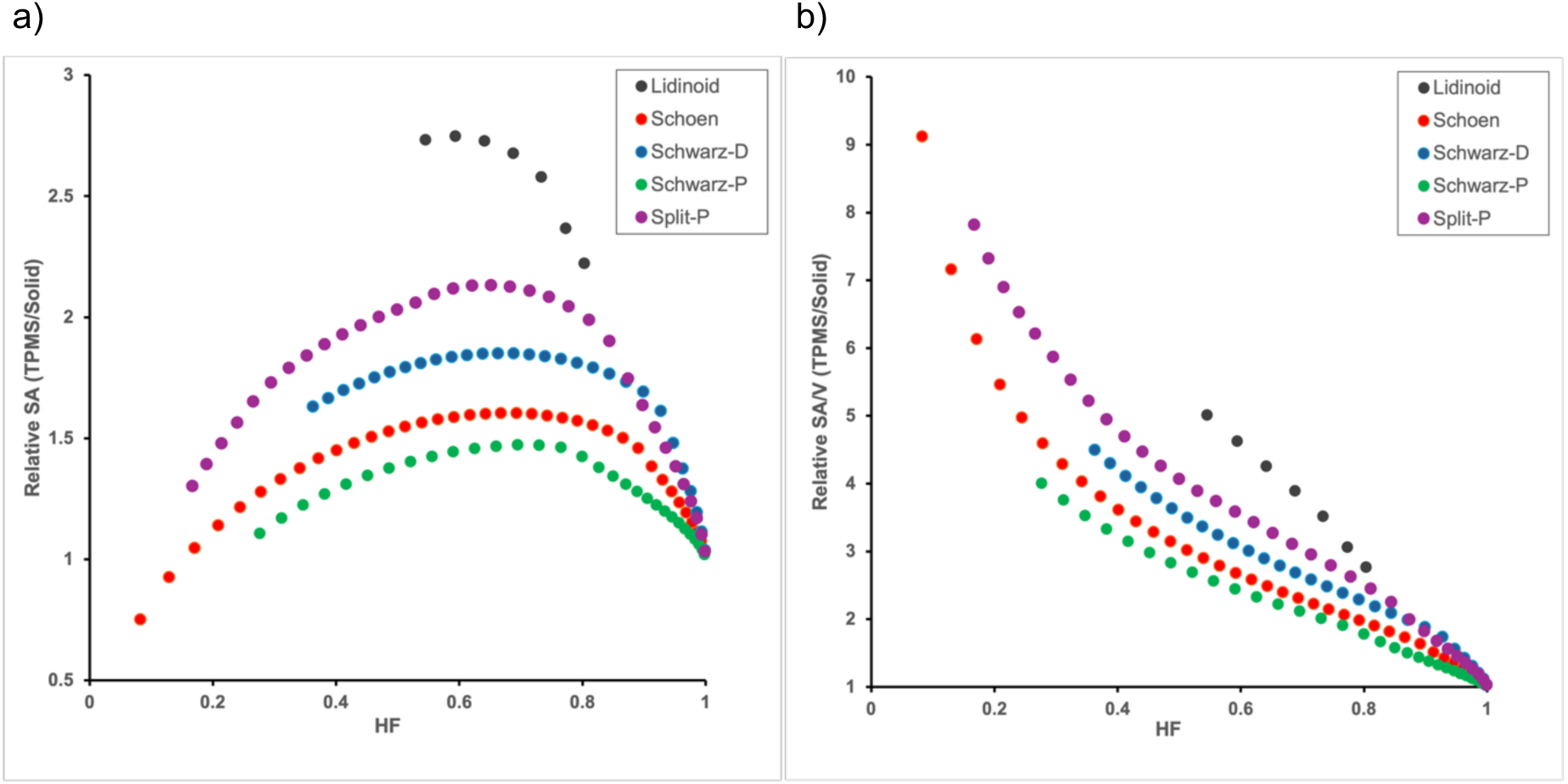
Geometric traits of the TPMS designs considered in this study. **a)** Relative surface area (SA), calculated as the SA of the TPMS design divided by the SA of a solid cube of the same dimensions, vs. relative hydrogel content (hydrogel fraction, HF) calculated as the hydrogel volume of the TPMS design divided by the hydrogel volume of a solid cube of the same dimensions. **b)** Relative surface-to-volume ratio (SA/V), calculated as the SA/V of the TPMS design divided by the SA/V of a solid cube of the same dimensions vs. the hydrogel fraction (HF). Only the realisations with HF ≈ 0.6 were 3D printed and tested with plants.

#### CAD models

A MATLAB code was developed to create the CAD models, with channels of diameters between 300 and 1,000 µm; all models were designed as 2×2×2 cm^3^ cubic structures, and their STL models are presented in Fig. 1. The MATLAB models were then converted into STL files compatible with the SLA printer used in this work. The surface area values reported throughout the manuscript correspond to the theoretical surface area calculated directly from the CAD models, and print fidelity was ensured by matching the minimum feature size of the designs to the printer resolution and verifying the dimensional accuracy of the printed substrates before experiments.

### Manufacturing

#### Substrate material

The following reagents were purchased from Sigma-Aldrich (Dorset, UK) unless stated otherwise: acrylamide (AAm; ≥ 99%), poly (ethylene glycol) diacrylate (PEGDA 700; average Mn 700), lithium phenyl-2,4,6-trimethylbenzoylphosphinate (LAP; ≥ 95%), 1-phenylazo-2-naphthol-6,8-disulfonic acid disodium salt (Orange G), Glycerol for molecular biology ≥99.0% (G5516), Murashige and Skoog (MS) plant growth medium MS Basal medium (M5519), MES hydrate (M8250) and Tris buffer (Fisher-Scientific, 10205100).

#### Photo-resin formulation

The water-based hydrogel formulation was developed for additive manufacturing (3D printing) using stereolithography (SLA), following a method we had previously established to consistently reproduce CAD-designed channels in the final substrate [20].

The photo-resin was prepared based on a final total mass of 100 g. In order of mixing, 5% (w/w) PEGDA_700_ cross-linker, 15% (w/w) AAm monomer and 5% (w/w) glycerol softening agent were added to a clean 150 ml amber bottle, where 80 g of deionised water was added and left to stir using a magnetic stirring plate for 30 min until fully dissolved. Next, relative to the total mass of monomer and cross-linker, 1% LAP and 0.24% Orange G were added to the mixture and left to stir for another 30 min until fully dissolved. The pH of the final photo-resin was adjusted to 5.7 using Tris buffer and 0.05% (w/v) MES hydrate prior to 3D printing.

#### 3D-printing

The final formulation was poured into a Formlabs Resin LT Tank and plugged into a Form 2 SLA printer (Formlabs, 405 nm light source, 140 µm laser spot size). ‘Open Mode’ setting was used which allows for use of 3rd party photo-resins. CAD models were uploaded using the Preform software with ‘Clear Resin V4’ settings, 100 µm layer height and no support structures. During the printing process, the print was paused every 10 layers to carefully remove any bubbles formed in the photo-resin due to the surface tension between the build plate and the tank. Bubbles were removed by gentle mixing of the photo-resin whilst the platform was raised under pause-mode.

Once the print was complete, the build platform was carefully detached from the printer, and printed substrates were immediately rinsed with DI water whilst attached to the platform to ensure excess photo-resin is removed. 3D-printed samples were then detached using a thin string and placed into a clear pot filled with DI water. The pots were gently agitated for 5 minutes and the process was repeated twice with fresh DI waterto ensure that excess photo-resin is removed within the network of channels of the hydrogels. Once fully cleaned, the DI water was replaced again, and the hydrogels were post-cured for 8 min using a FormCure (Formlabs) post-curing station with a 405 nm light source to photo-cure any unreacted monomers and cross-linkers.

#### Dehydration and rehydration

The 3D-printed hydrogels were dehydrated in open air at room temperature for 24 hours. Once dehydrated, the hydrogels were then soaked in ¼ Murashige and Skoog (MS) plant growth medium: 0.1075% (w/v) MS basal medium, 0.05% (w/v) MES hydrate, pH adjusted to 5.7 with Tris buffer to allow the hydrogels to absorb the nutrients necessary for plant growth. The soaking medium was discarded and replaced with fresh ¼ MS medium daily for 7 days, to remove any remaining monomers and cross-linkers from the hydrogel that may have not post-cured.

The hydrogel growth substrates are non-toxic following the process of washing, post-curing, hydration and dehydration, which ensures unreacted chemical components are either photo-cured and/or washed out. Further, after dehydration and hydration, the hydrogels have a buffer pH of 5.7, and contain sufficient micronutrients to support the germination of Arabidopsis seeds placed on their surface and the subsequent plant growth to flowering.

The mechanical and chemical properties of our hydrogel system have been previously optimised for high-resolution 3D-printing in our previous work [20].

### Plant germination

#### Hydrogel

Wild-type *Arabidopsis thaliana* seeds, Columbia (Col-0) ecotype, were used for this work. Seeds were imbibed in water and kept in the dark at 4°C for two days to synchronise germination. Seeds were surface sterilised with 50% (v/v) Haychlor bleach and 0.0005% (v/v) Triton X-100 for 3 min and then rinsed with sterilised Milli-Q water six times. Sterilised seeds were sown on the top surface of each hydrogel print, which were then placed inside transparent Magenta boxes GA-7-3 77×77×77 mm^3^ (Sigma-Aldrich, V8380). The boxes were transferred to a growth chamber at 22 °C, with a (16 h : 8 h) = (light : dark) photoperiod and light intensity 120 µmol m^−2^ s^−1^.

#### Hydroponics

For the hydroponics control, seeds were sown in PCR tubes containing ¼ MS solid medium (0.1075% (w/v) MS Basal medium, 0.05% (w/v) MES hydrate, 0.8% (w/v) agar (Sigma-Aldrich 05040), pH adjusted to 5.7 with Tris Buffer. The tips of the PCR tubes were cut to allow the growing roots to come out, and then were inserted into a 3D-printed support placed inside Magenta boxes (Sigma-Aldrich, V8380) filled with ¼ MS liquid medium, as previously described [29]. A 0.5 µm diffusion stone connected to an air pump was placed inside each Magenta box, to aerate the liquid medium. The Magenta boxes were then transferred to a growth chamber at 22 °C, with a (16 h : 8 h) = (light : dark) photoperiod and light intensity 120 µmol m^−2^ s^−1^.

### Experimental design

For the hydrogel prints, two sterilised *Arabidopsis* seeds were sown on top of each print, and two prints were placed in each Magenta box; five Magenta boxes were used for each hydrogel design. For the hydroponics control, one sterilised seed was sown in each PCR tube, and 10 tubes were placed in each Magenta box filled with liquid growth medium; five Magenta boxes were used in hydroponic conditions.

Plantlets were left on the same substrate and grown for 1 to 5 weeks after sowing. One control was constituted by a “solid” substrate, 3D-printed with the same hydrogel composition but without channels. An aerated liquid culture (hydroponics) containing the same concentration of micronutrients used in the hydrogel was adopted as a second control representing the traditional hydroponic cultivation currently mostly popular in soilless indoor farming.

The entire pipeline, from 3D printing to phenotyping, is summarised in Supplementary Fig. 1.

### Phenotyping

At the end of each week, leaves on each plant were counted and cut from the plants with dissecting scissors (Wolflabs 503667) and laid flat on the surface of a ¼ MS agar plate. The leaves were then visualised under a stereo microscope (Nikon SMZ800N), and images were recorded with a Nikon DS-Fi3 camera using Nikon NIS-Elements Imaging software.

The area of the leaves was determined using Image-J software. For each image, a colour intensity threshold was manually determined, which was used to generate a contour around the leaf area. The threshold was adjusted as needed for the contour to best represent the area. The total leaf area was then calculated and recorded from the number of pixels within the contour. The flowering time and the number of flowering plants were monitored throughout the experiments without disrupting plants’ growth, through a viewing window on the Magenta box lids modified with a custom-cut glass slide (VWR MARI1100420).

### Statistical analysis

The statistical analysis on leaf number and area data was done by comparing the five different hydrogel designs first, and then the hydroponics control versus each hydrogel design. The normality of the samples was checked with the Shapiro-Wilk test with alpha-level = 0.05. Levene’s test with alpha-level = 0.05 was used to assess the equality of variances. For the comparisons among the hydrogel designs, if the distributions were normally distributed and the variances were equal, one-way ANOVA at the 0.05 level with Tukey’s multiple comparison test was used. For non-normally distributed data and/or unequal variances among the groups, Kruskal-Wallis ANOVA at the 0.05 level with Dunn’s test was performed. For the comparison between each hydrogel design and the hydroponics control, when the samples were normally distributed, the two-sample *t*-test with Bonferroni’s correction was used. If one of the distributions was not normal, the Mann-Whitney test with Bonferroni’s correction was used. Following Bonferroni-s correction, the alpha level for the two-sample *t*-test and Mann-Whitney test = 0.008. The association between flowering proportion and relative surface area was assessed by Pearson correlation; significance was determined by a two-tailed t-test (n = 5, df = 3).

All statistical tests were performed with Origin Pro.

In the figures, \**P* < 0.05, \*\**P* < 0.01, and \*\*\**P* < 0.001 in comparisons among hydrogel designs, \**P* < 0.008, \*\**P* < 0.002, and \*\*\**P* < 0.0002 in comparison between hydroponics vs hydrogel designs.

## RESULTS

### Arabidopsis germination and growth on the 3D-printed hydrogel substrates

The five TPMS geometries were successfully 3D-printed as self-supporting hydrogel substrates (Fig. 1a,b). *Arabidopsis* seeds sown on their top surface germinated and grew (Fig. 1c), and electron microscopy confirmed that the plant roots grow on the inner surface of the channels (Supplementary Fig. 2). To compare the effectiveness of the network designs among themselves and with traditional hydroponics, we then quantified three developmental traits: leaf number, leaf area and flowering time.

### Leaf number

To quantify vegetative growth, we focused on leaf development. After the first week, the two cotyledons (embryonic leaves) emerged from all the plants grown in the five substrates and the hydroponics (Fig. 3 a). A few plants grew true leaves in the Lidinoid, but also in the solid substrate and hydroponics; plants grown in the Lidinoid substrate developed an average of 2.8 leaves per plant, showing the best yield among all the designs (Fig. 3 a, Dunn’s test between Lidinoid and Schoen, Lidinoid and Schwarz-D, Lidinoid and Schwarz-P, Lidinoid and Split-P, *P*=0.018).

**Fig. 3.**
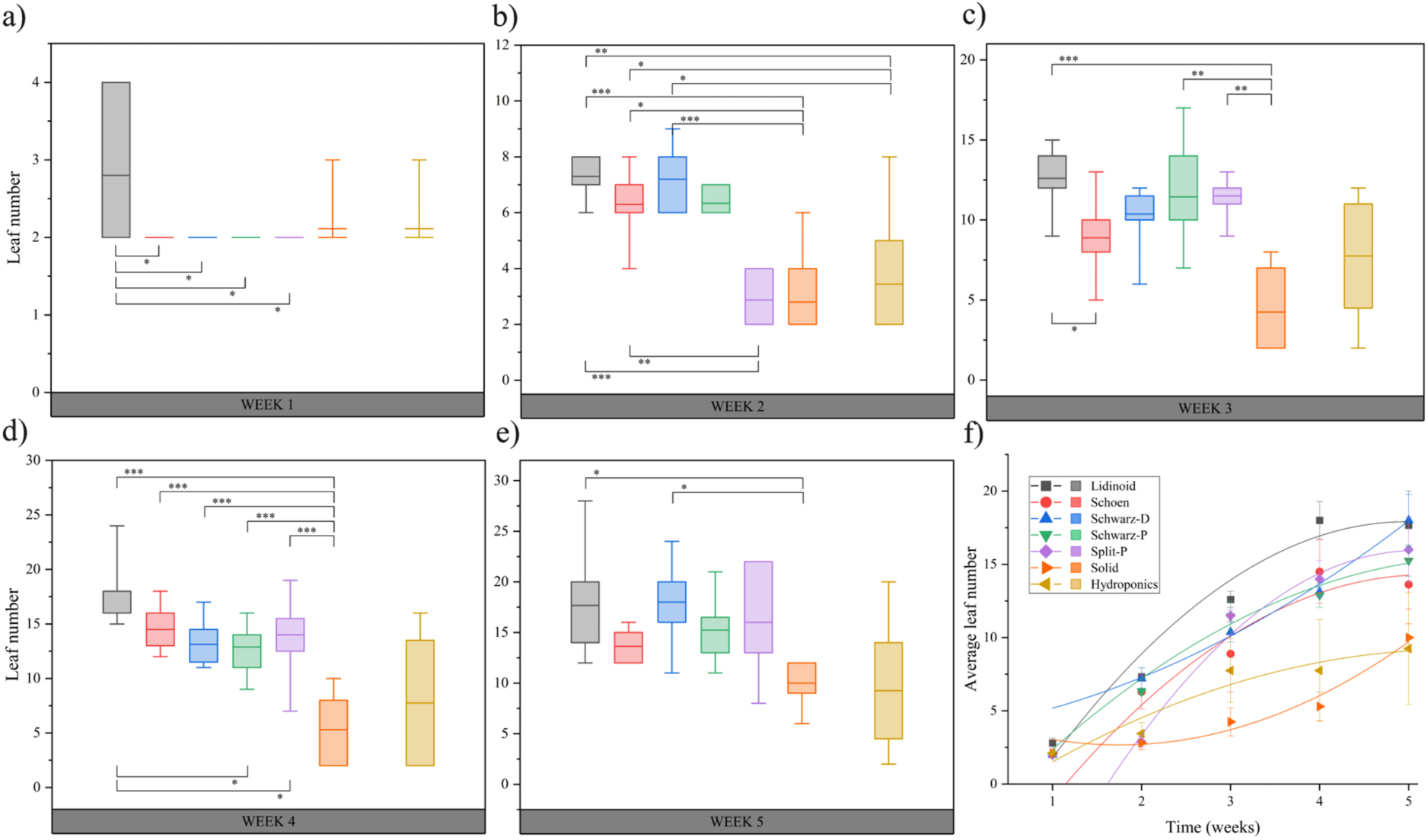
Average number of leaves per plant grown in each substrate design (the five TPMS designs realised at HF ≈ 0.6), with solid (HF = 1) and hydroponics as controls. **a - e)** Each box-plot represents the distribution of these averages across the population of plants grown in that substrate; the lower and upper limits of each box represent the 25th and 75th percentiles of the data, respectively, the whiskers represent the 5^th^ and 95th percentiles, and the horizontal line represents the mean. **f)** Each symbol represents the average leaf number of all plants grown in each substrate, at the corresponding week. Error bars, s.e.m.

After 2 weeks, plants grown in Split-P, solid, and hydroponics still had only the two cotyledons, while all the other designs developed non-embryonic leaves as well (Fig. 3 b). Among these, plants on Lidinoid, Schwarz-D and Schoen were more efficient in leaf number development than the ones on solid hydrogel (Fig. 3 b, Dunn’s test between solid and Lidinoid, and solid and Schwarz-D, *P*<0.001; Dunn’s test between solid and Schoen, *P*<0.05) and the hydroponics control (Mann-Whitney test with Bonferroni correction between hydroponics and Lidinoid, *P*= 0.001, hydroponics vs Schoen, and hydroponics vs Schwarz-D, *P*<0.008).

In the subsequent weeks, the number of leaves developed by plants grown in the Lidinoid substrate was always greater than those grown in the solid hydrogel (see Supplementary Table 1 for detailed statistics). After four weeks, Lidinoid, Schoen, Schwarz-D, Schwarz-P and Split-P all generated more leaves than the solid hydrogel (Fig. 3 d, Tukey test, solid vs Lidinoid, solid vs Schoen, solid vs Schwarz-D, solid vs Schwarz-P, solid vs Split-P, *P*<0.001).

A statistically significant difference between the number of leaves developed in any patterned hydrogel and hydroponics was observed only at week 2 (Fig. 3 b; see Supplementary Table 2 for detailed statistics). Notably, plants grown in hydroponics showed unusually large variability in leaf number (Fig. 3 c, d, and e). Moreover, when plotting the number of emerged leaves as a function of time (Fig. 3 f), it appears that the rate of leaf production was higher for plants grown in patterned hydrogels, compared to those grown in solid hydrogel or hydroponics.

### Leaf area

To further quantify the effect of patterned hydrogel substrate on shoot growth, we also measured the average leaf size in each plant and compared those grown on patterned substrates to those grown in solid hydrogel or hydroponics. Briefly, all leaves were cut from each plant weekly, and their area was individually measured with the aid of a dissecting microscope and Image-J (see Methods).

After only one week, Lidinoid was the patterned substrate where plants developed the largest average leaf size (Fig. 4 a, see Supplementary Tables 3 and 4 for detailed statistics). Moreover, in the first week, plants in all patterned hydrogels showed an average leaf area larger than that of the hydroponics control (Fig. 4 a, see Supplementary Table 4 for detailed statistics).

**Fig. 4.**
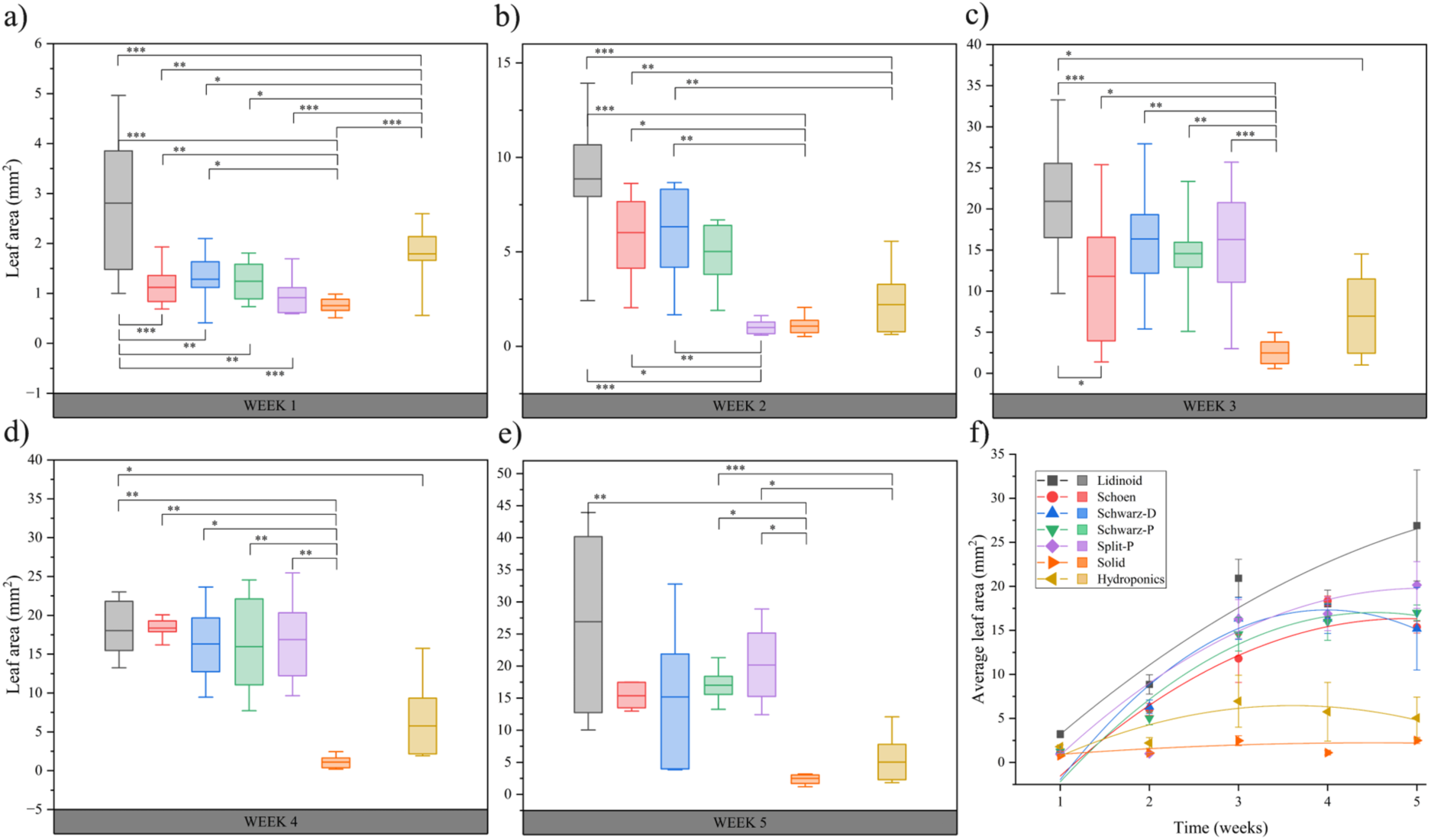
Average leaf area per plant grown in each substrate design (the five TPMS designs realised at HF ≈ 0.6), with solid (HF = 1) and hydroponics as controls. **a - e)** Each box-plot represents the distribution of these averages across the population of plants grown in that substrate; the lower and upper limits of each box represent the 25th and 75th percentiles of the data, respectively, the whiskers represent the 5^th^ and 95th percentiles, and the horizontal line represents the mean. **f)** Each symbol represents the average leaf area of all plants grown in each substrate, at the corresponding week. Error bars, s.e.m.

After two weeks, plants grown in Lidinoid, Schoen, and Schwarz-D developed leaves bigger than those grown on the solid hydrogel (Fig. 4 b, Dunn’s test between solid and Lidinoid, *P*<0.001, solid and Schoen, *P*=0.014, solid and Schwarz-D, *P*<0.01) and the hydroponics control (Fig. 4 b, two-sample *t*-test with Bonferroni correction between hydroponics and Lidinoid, *P*<0.0002, hydroponics vs Schoen and hydroponics vs Schwarz-D, *P*<0.002).

After three and four weeks, all the plants on patterned hydrogels on average had larger leaves than those on the solid hydrogel (Fig. 4 c, and d, see Supplementary Table 3 for detailed statistics), corroborating the results on leaf number at week 4 (Fig. 3 d). During the first four weeks, the Lidinoid design performed consistently better in leaf size compared to the traditional hydroponics (Fig. 4 a-d; see Supplementary Table 3 for detailed statistics). However, the statistical significance was lost during week 5, coinciding with substantial variability among plants grown in the Lidinoid (Fig. 4 e). In fact, after five weeks, the three plants that developed the biggest average leaf area of all (38.3, 40.2, and 43.9 mm^2^) were all grown in the Lidinoid hydrogels (Fig. 4 e). Furthermore, the plot of average leaf area as a function of time (Fig. 4 f) indicates that the plants growing on the Lidinoid were the ones with the highest rate of leaf growth, while the ones growing in solid hydrogel or hydroponics were the ones with the lowest rate.

Finally, except for week 1 (Fig. 4 a, two-sample *t*-test with Bonferroni correction between hydroponics and solid, *P*<0.0002), the solid hydrogel and the hydroponics substrates were always indistinguishable from each other both in leaf number and average leaf area (see Supplementary Tables 1 - 4 for detailed statistics).

### Flowering

Besides assessing vegetative growth with leaf development, we also analysed reproductive growth by quantifying the time required to develop the first flowers, or flowering time. Strikingly, plants grown in Lidinoid, Schoen, Schwarz-D and Split-P patterned hydrogels produced flowers within five weeks (35 days) after sowing, while plants grown in Schwarz-P, solid hydrogels and hydroponics did not (Fig. 5). The first flowering event took place in the Lidinoid hydrogel on day 25, followed by Split-P on day 26 and Schoen and Schwarz-D on day 28 (Fig. 5). By the end of the five weeks, 100% (± 13.4% s.e.p.) of the plants grown in Lidinoid flowered, while only 60% (± 9.5% s.e.p.) of plants in Split-P, 30% (± 12.7% s.e.p.) in Schwarz-D and 10% (± 9.5% s.e.p.) in Schoen hydrogels flowered (Fig. 5). Plants in the Lidinoid hydrogel were the fastest to reach the vegetative-reproductive developmental transition, with all plants flowering after only four weeks (Fig. 5).

**Fig. 5.**
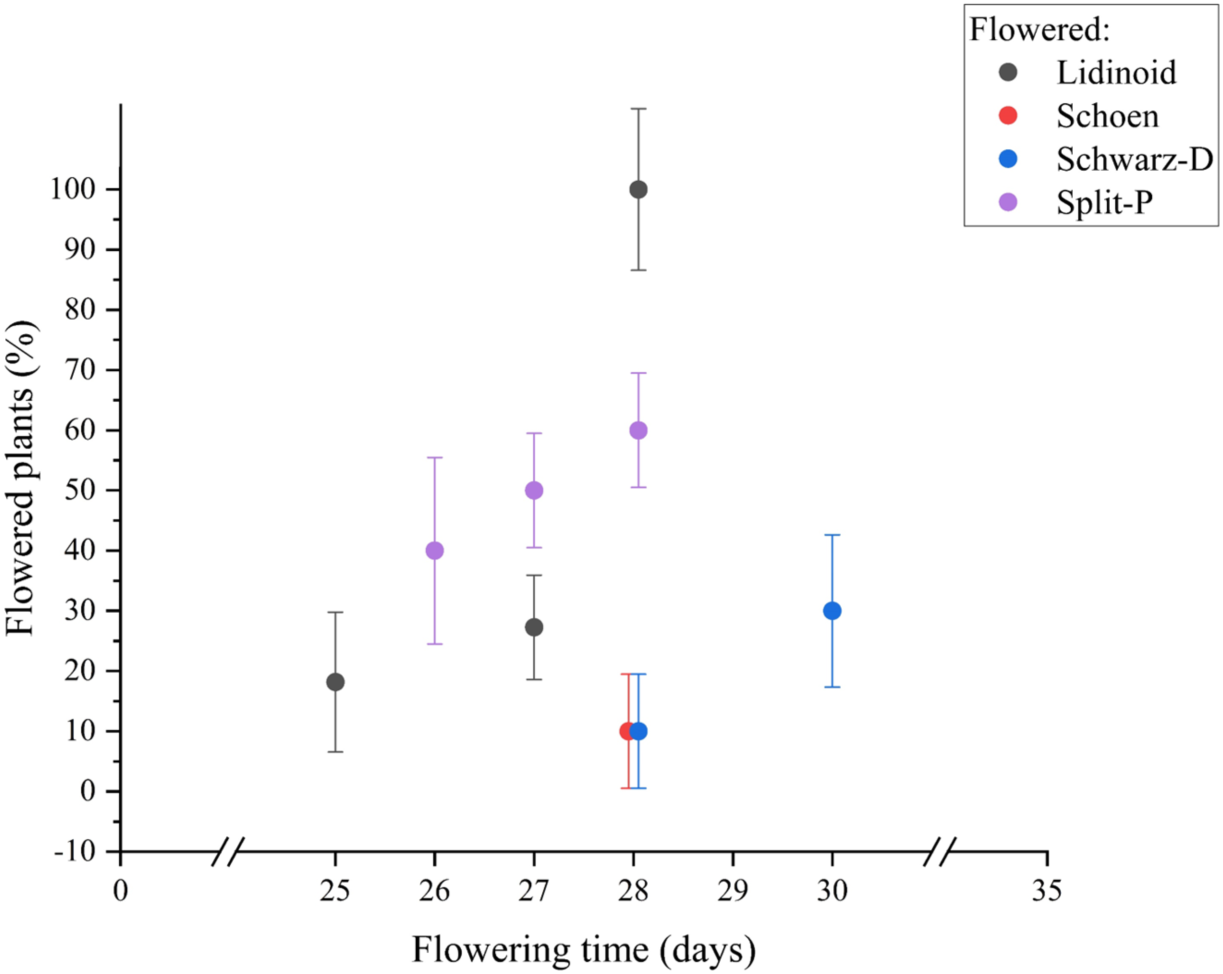
Percentage of plants flowering each day, when grown in each substrate design (the five TPMS designs realised at HF ≈ 0.6), with solid (HF = 1) and hydroponics as controls. Plants were monitored for a total of five weeks (35 days). Plants grown in solid hydrogel, hydroponics and Schwarz-P did not flower in the first 35 days. Error bars, s.e.p.

Interestingly, when considering the total percentage of flowered plants, the performance of each TPMS design is positively correlated to the relative surface area (Fig. 6; Pearson r = 0.996, R² = 0.991, n = 5, p-value = 3.6 × 10⁻⁴), i.e. with a higher surface area corresponding to a higher % of flowering plants at day 35.

**Fig. 6.**
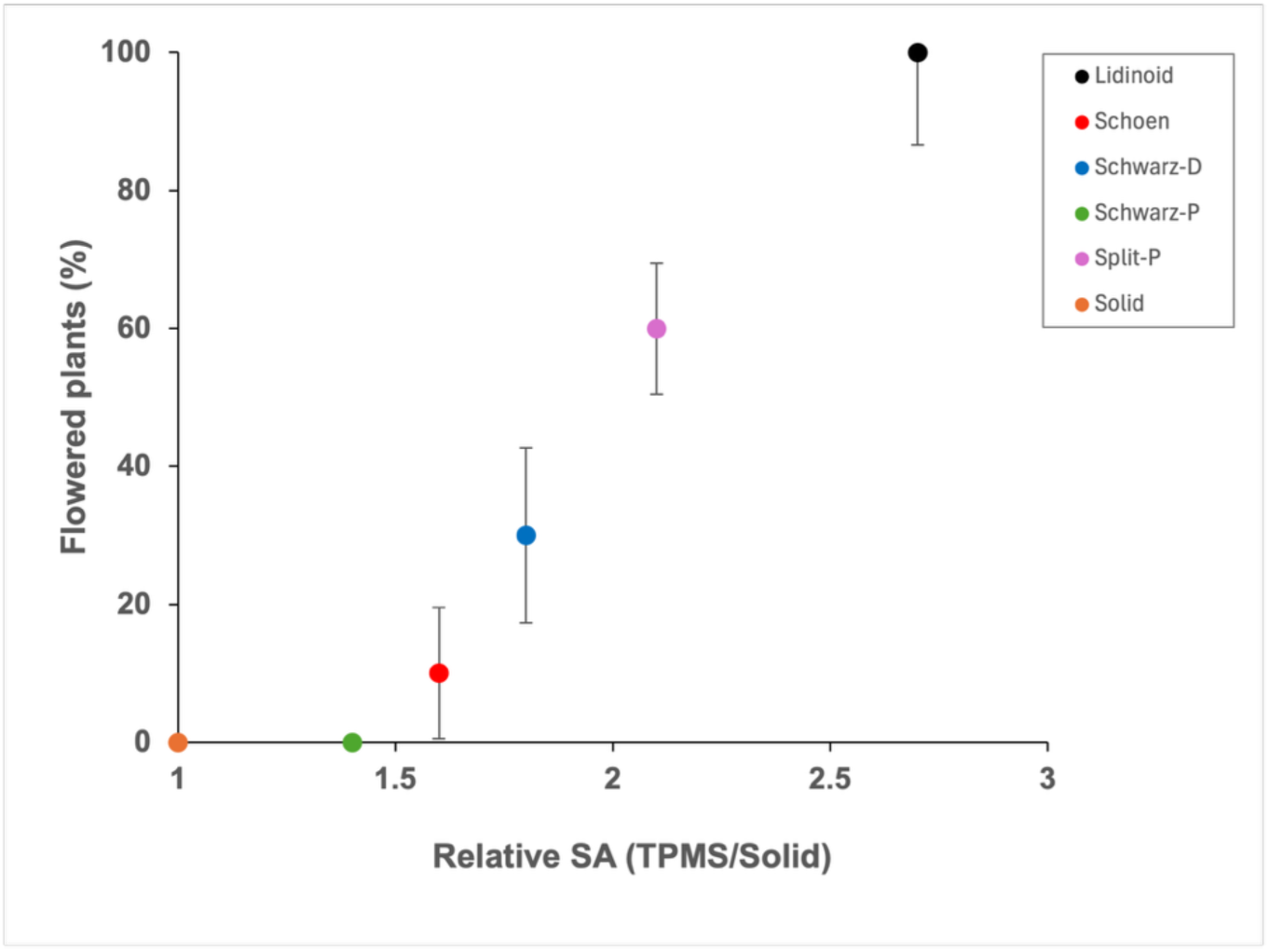
Correlation between the proportion of flowering plants at day 35 and the relative surface area (SA) of each substrate design (the five TPMS designs realised at HF ≈ 0.6), with solid (HF = 1) as control. SA was calculated as the surface of the TPMS design divided by the surface of a solid cube of the same dimensions.

## DISCUSSION

The idea of 3D printing an artificial substrate to replicate some of the physical and topological traits of natural soil is not entirely new [21–23]. Previous efforts to engineer artificial soil analogues have demonstrated the feasibility of using porous materials and additively manufactured architectures to influence root growth and water transport. However, most existing approaches have focused on reproducing bulk mechanical properties or generating stochastic pore networks, for example, using X-ray computed tomography to image undisturbed soil samples and subsequently 3D-print replicas of natural pore geometry [22,23]. More recent work has explored 3D-printed hydrogels with simple orthogonal channel lattices for root development [21], yet without a systematic comparison of alternative geometries. A key limitation of these studies is that they do not vary internal topology in a controlled manner, making it difficult to determine which geometric parameters, if any, are critical for plant growth.

What remains less clear is which of these traits suffices to support the long-term and robust growth of an entire plant. More specifically, what is the minimal design and composition required to create an artificial substrate on which a flowering plant can germinate and grow leaves and flowers?

In contrast, our study here compares five distinct TPMS architectures, each with a well-defined surface-to-volume ratio at a nearly identical hydrogel fraction (HF ≈ 0.6), enabling direct assessment of how channel geometry influences vegetative growth and the transition to flowering. The use of high-resolution stereolithography enables the fabrication of these complex, fully interconnected networks that remain continuously exposed to the atmosphere, a capability difficult to achieve with conventional manufacturing methods.

In this work, we show that a block of hydrogel, patterned with an internal periodic network of air-filled tubules, is sufficient to sustain germination, vegetative growth, and flowering in the model plant *Arabidopsis thaliana*. Moreover, we show that the specific geometry of the periodic tubule network significantly influences plant growth.

Among the TPMS patterns tested, the Lidinoid design outperformed the others in terms of average number (Fig. 3) and size (Fig. 4) of leaves, as well as flowering time (Fig. 5). Notably, this design offers the highest surface-to-volume ratio among all tested geometries, given the near-identical hydrogel volume fraction across designs. This outcome is perhaps expected, as a higher surface-to-volume ratio may improve gas exchange between the water in the hydrogel and the air within the tubule network. One limitation of hydroponic (i.e. liquid-based) cultivation is the relatively low concentration of dissolved oxygen in water, which can constrain root development [30]. Roots require oxygen for cellular respiration, particularly during active growth phases supporting leaf and flower development. The 3D printed substrate described here can be thought of as a “vascularised”, or patterned, hydrogel, with the tubule network that we propose facilitates passive diffusion and enhances oxygen availability.

A higher relative surface area increases the interface across which roots exchange gases, water and nutrients with the hydrogel, so the correlation is consistent with enhanced oxygen availability but equally with improved water and nutrient exchange or greater root–substrate contact; none of these was directly measured. Of these, the contribution of contact area is likely limited, since our channel diameters (300–1,000 µm) are considerably larger than typical *Arabidopsis* root diameters (∼100 µm). Substrate properties not tied to the interface area, such as altered mechanical properties affecting root growth, may also contribute. Furthermore, relative surface area may correlate with other geometric properties of the channel networks, such as tortuosity and pore connectivity, which could independently influence root growth and warrant future investigation. Disentangling these contributions will require future direct oxygen measurements and targeted perturbation experiments.

One of the more striking results in this work is that the patterned hydrogel outperforms a traditional hydroponic substrate in promoting leaf number and size (Fig. 3 and 4), particularly when using the Lidinoid design. This is further supported by the observation that an unpatterned hydrogel block performs comparably to hydroponics, indicating that it is the geometry of the internal network, rather than the hydrogel material itself, that drives the difference.

Finally, we argue that quantitative data on flowering provide a critical metric for evaluating substrate functionality. In *Arabidopsis,* the ability to transition efficiently from vegetative growth to flowering is a key determinant of plant fitness [31] and has direct implications for fruit production, an important consideration in agritech applications such as indoor crop cultivation. Notably, the Lidinoid design not only produced the fastest leaf growth rate among all tested geometries (Fig. 4f), but also promoted the earliest flowering (Fig. 5), suggesting that accelerated vegetative development translates into earlier reproductive transition. The monotonic increase in flowering proportion with relative surface area across the five designs (Fig. 6) indicates that performance scales with the *extent* of the internal hydrogel interface, not merely with the presence of channels. This identifies relative surface area as a dominant geometric parameter, although still leaving open which of the interface-dependent processes is responsible.

From an application perspective, additive manufacturing remains slower and more costly than conventional substrate production, currently limiting practical use to research laboratories and controlled-environment agriculture facilities with access to high-resolution 3D printing. This study represents a proof-of-principle validated under controlled conditions (TRL 4), still far from commercial readiness; future work must evaluate performance with crop species under realistic cultivation conditions. The design was optimised for direct seed germination, and further development is required to accommodate transplant-based cultivation and vegetative propagation.

Taken together, our findings support a broader strategy of using additive manufacturing (3D printing) as a powerful method for fabricating finely patterned hydrogels for indoor plant cultivation. A simple, though certainly not exhaustive, explanation is that the hydrogel provides water and nutrients, while the interconnected channels could enable passive oxygen delivery to root meristems. In this respect, the approach echoes the way air-filled pores aerate roots in natural soil, although a direct comparison with soil lies beyond the scope of this study.

## ACKOWLEDGEMENTS

We would like to thank Dr Borut Lampret for helping with code development and analysis tools, Dr Livia Kalossaka for their assistance during the early development of this study.

---

**Supplementary Fig. 1.**
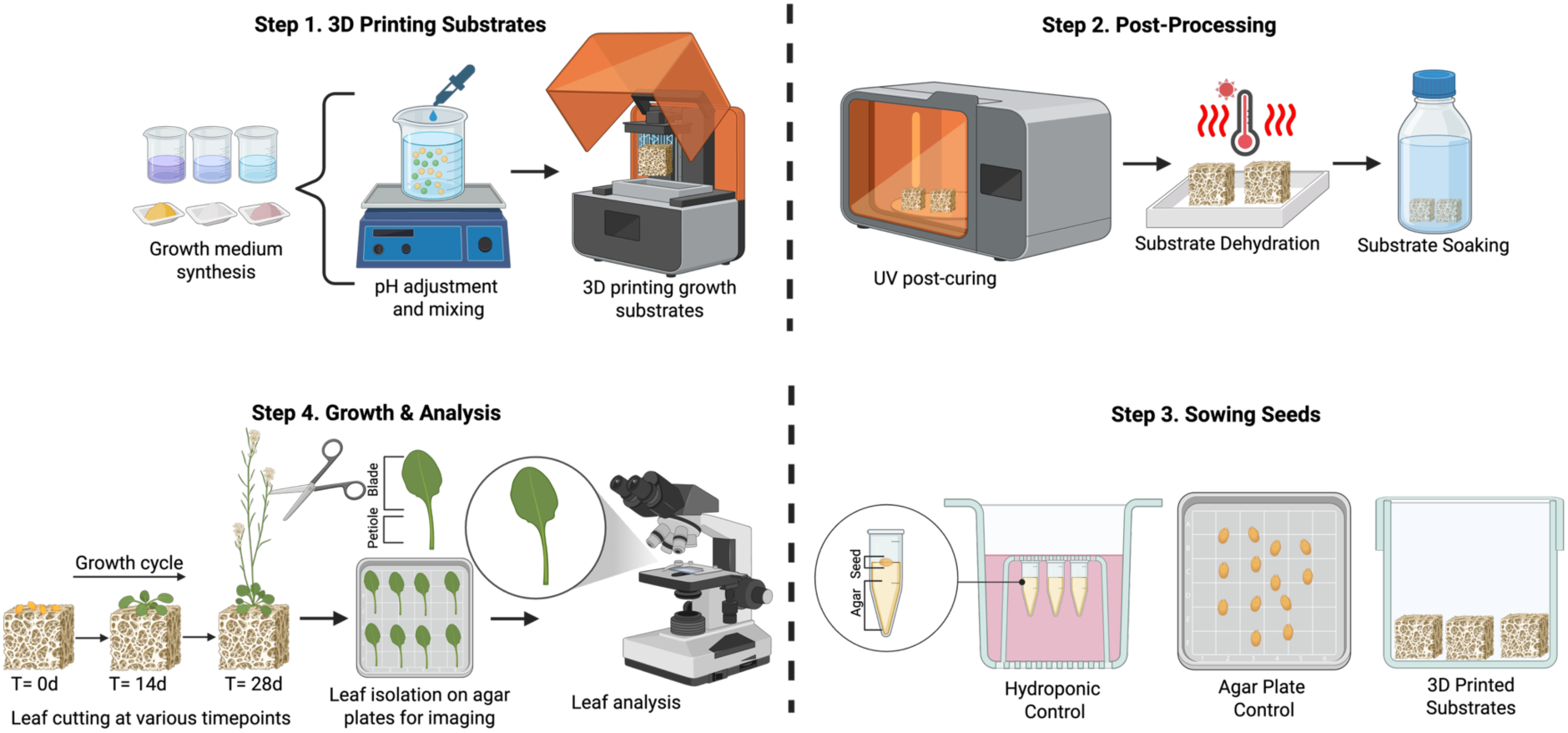
Schematic of the experimental pipeline used in this work including 3D printing, post-processing, sowing seeds, and growth and analysis.

**Supplementary Fig. 2.**
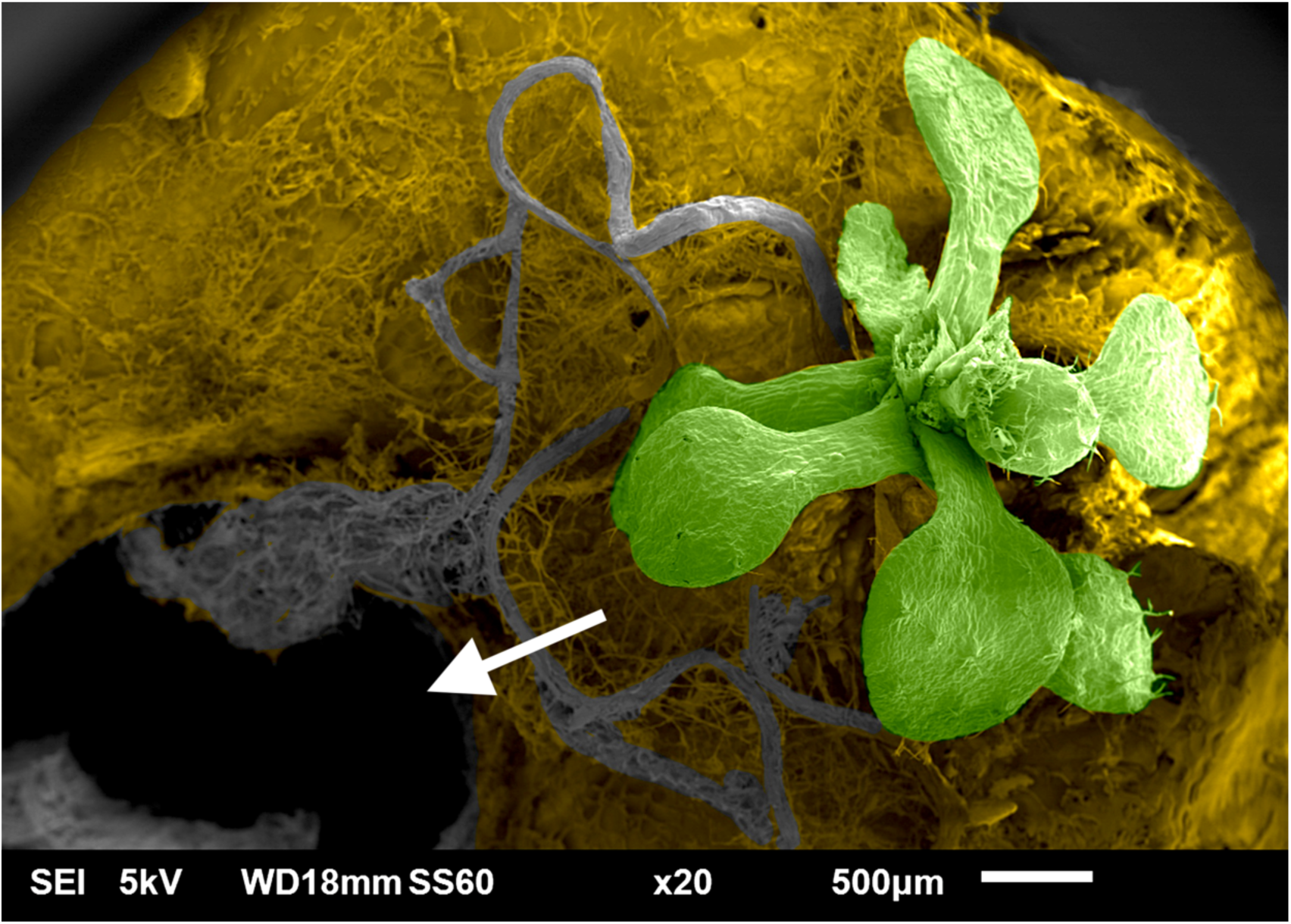
Scanning Electron Microscopy (SEM) image of an Arabidopsis seedling with its primary root entering one of the channels (white arrow) of the 3D-printed hydrogel. False-coloured: primary root, grey; shoot, green; hydrogel, yellow. Scale bar, 500µm.

**Supplementary Table 1.** Statistical analysis of leaf number among the five hydrogel TPMS designs plus the solid control. Statistically significant differences in bold.

|  | WEEK 1 |  | WEEK 2 |  | WEEK 3 |  | WEEK 4 |  | WEEK 5 |  |
| --- | --- | --- | --- | --- | --- | --- | --- | --- | --- | --- |
|  | Kruskal-Wallis ANOVA | Dunn's test | Kruskal-Wallis ANOVA | Dunn's test | Kruskal-Wallis ANOVA | Dunn's test | One-way ANOVA | Tukey's test | One-way ANOVA | Tukey's test |
|  | *P*-value | *P*-value | *P*-value | *P*-value | *P*-value | *P*-value | *P*-value | *P*-value | *P*-value | *P*-value |
| Overall | **0.005** |  | **4E-07** |  | **4E-05** |  | **1E-09** |  | **0.012** |  |
| Schoen vs lidinoid |  | **0.018** |  | 1 |  | **0.037** |  | 0.288 |  | 0.412 |
| Schwarz-D vs lidinoid |  | **0.018** |  | 1 |  | 0.824 |  | **0.031** |  | 1 |
| Schwarz-D vs schoen |  | 1 |  | 1 |  | 1 |  | 0.945 |  | 0.326 |
| Schwarz-P vs lidinoid |  | **0.018** |  | 1 |  | 1 |  | **0.017** |  | 0.860 |
| Schwarz-P vs schoen |  | 1 |  | 1 |  | 1 |  | 0.887 |  | 0.960 |
| Schwarz-P vs schwarz-D |  | 1 |  | 1 |  | 1 |  | 1 |  | 0.782 |
| Solid vs lidinoid |  | 0.284 |  | **8E-05** |  | **3E-05** |  | **0** |  | **0.020** |
| Solid vs schoen |  | 1 |  | **0.025** |  | 1 |  | **3E-06** |  | 0.532 |
| Solid vs schwarz-D |  | 1 |  | **3E-04** |  | 0.105 |  | **1E-05** |  | **0.014** |
| Solid vs schwarz-P |  | 1 |  | 0.164 |  | **0.009** |  | **1E-05** |  | 0.157 |
| Split-P vs lidinoid |  | **0.018** |  | **4E-04** |  | 1 |  | 0.117 |  | 0.976 |
| Split-P vs schoen |  | 1 |  | 0.055 |  | 0.653 |  | 1 |  | 0.868 |
| Split-P vs schwarz-D |  | 1 |  | **0.001** |  | 1 |  | 0.990 |  | 0.948 |
| Split-P vs schwarz-P |  | 1 |  | 0.257 |  | 1 |  | 0.965 |  | 1 |
| Split-P vs solid |  | 1 |  | 1 |  | **0.002** |  | **1E-06** |  | 0.111 |

**Supplementary Table 2.** Statistical analysis of leaf number between the hydroponics control and the five hydrogel TPMS designs plus the solid control. Bonferroni’s correction was applied to all tests. Statistically significant differences in bold.

|  | WEEK 1 | WEEK 2 | WEEK 3 | WEEK 4 | WEEK 5 |
| --- | --- | --- | --- | --- | --- |
|  | Mann-Whitney test | Mann-Whitney test | Mann-Whitney test | Two-sample t-test | Two-sample t-test |
|  | *P*-value | *P*-value | *P*-value | *P*-value | *P*-value |
| Lidinoid vs hydroponics | 0.108 | **0.001** | 0.022 | 0.012 | 0.081 |
| Schoen vs hydroponics | 0.947 | **0.007** | 0.797 | 0.145 | 0.336 |
| Schwarz-D vs hydroponics | 0.947 | **0.002** | 0.364 | 0.220 | 0.048 |
| Schwarz-P vs hydroponics | 0.947 | 0.014 | 0.210 | 0.237 | 0.215 |
| Solid vs hydroponics | 1 | 0.628 | 0.190 | 0.364 | 0.860 |
| Split-P vs hydroponics | 0.947 | 0.984 | 0.074 | 0.061 | 0.139 |

**Supplementary Table 3.** Statistical analysis of the average leaf area among the five hydrogel TPMS designs plus the solid control. Statistically significant differences in bold.

|  | WEEK 1 |  | WEEK 2 |  | WEEK 3 |  | WEEK 4 |  | WEEK 5 |  |
| --- | --- | --- | --- | --- | --- | --- | --- | --- | --- | --- |
|  | Kruskal-Wallis ANOVA | Dunn's test | Kruskal-Wallis ANOVA | Dunn's test | One-way ANOVA | Tukey's test | Kruskal-Wallis ANOVA | Dunn's test | Kruskal-Wallis ANOVA | Dunn's test |
|  | *P*-value | *P*-value | *P*-value | *P*-value | *P*-value | *P*-value | *P*-value | *P*-value | *P*-value | *P*-value |
| Overall | **9E-12** |  | **2E-07** |  | **1E-05** |  | **2E-04** |  | **0.005** |  |
| Schoen vs lidinoid |  | **8E-05** |  | 1 |  | **0.037** |  | 1 |  | 1 |
| Schwarz-D vs lidinoid |  | **0.004** |  | 1 |  | 0.665 |  | 1 |  | 1 |
| Schwarz-D vs schoen |  | 1 |  | 1 |  | 0.695 |  | 1 |  | 1 |
| Schwarz-P vs lidinoid |  | **0.003** |  | 0.792 |  | 0.280 |  | 1 |  | 1 |
| Schwarz-P vs schoen |  | 1 |  | 1 |  | 0.941 |  | 1 |  | 1 |
| Schwarz-P vs schwarz-D |  | 1 |  | 1 |  | 0.993 |  | 1 |  | 1 |
| Solid vs lidinoid |  | **4E-11** |  | **1E-05** |  | **3E-06** |  | **0.005** |  | **0.007** |
| Solid vs schoen |  | 0.156 |  | **0.014** |  | **0.048** |  | **0.002** |  | 0.159 |
| Solid vs schwarz-D |  | **0.008** |  | **0.005** |  | **0.001** |  | **0.010** |  | 0.170 |
| Solid vs schwarz-P |  | **0.011** |  | 0.284 |  | **0.004** |  | **0.008** |  | **0.016** |
| Split-P vs lidinoid |  | **9E-09** |  | **2E-05** |  | 0.594 |  | 1 |  | 1 |
| Split-P vs schoen |  | 1 |  | **0.017** |  | 0.655 |  | 1 |  | 1 |
| Split-P vs schwarz-D |  | 0.152 |  | **0.007** |  | 1 |  | 1 |  | 1 |
| Split-P vs schwarz-P |  | 0.198 |  | 0.275 |  | 0.992 |  | 1 |  | 1 |
| Split-P vs solid |  | 1 |  | 1 |  | **5E-04** |  | **0.007** |  | **0.014** |

**Supplementary Table 4.** Statistical analysis of the average leaf area between hydroponics control and the five hydrogel TPMS designs plus the solid control. Bonferroni’s correction was applied to all tests. Statistically significant differences in bold.

|  | WEEK 1 | WEEK 2 | WEEK 3 | WEEK 4 | WEEK 5 |
| --- | --- | --- | --- | --- | --- |
|  | Two-sample *t*-test | Two-sample *t*-test | Two-sample *t*-test | Two-sample *t*-test | Two-sample *t*-test |
|  | *P*-value | *P*-value | *P*-value | *P*-value | *P*-value |
| Lidinoid vs hydroponics | **1E-04** | **8E-05** | **0.004** | **0.005** | 0.027 |
| Schoen vs hydroponics | **5E-04** | **7E-04** | 0.311 | 0.031 | 0.018 |
| Schwarz-D vs hydroponics | **0.007** | **7E-04** | 0.039 | 0.009 | 0.138 |
| Schwarz-P vs hydroponics | **0.004** | 0.012 | 0.051 | 0.023 | **2E-04** |
| Solid vs hydroponics | **2E-06** | 0.104 | 0.226 | 0.258 | 0.362 |
| Split-P vs hydroponics | **2E-05** | 0.085 | 0.034 | 0.011 | **0.004** |

## REFERENCES

1. Ben-Noah I, Friedman SP. Review and Evaluation of Root Respiration and of Natural and Agricultural Processes of Soil Aeration. Vadose Zone J. 2018;17:1–47. 10.2136/vzj2017.06.0119

2. Amthor JS. The McCree–de Wit–Penning de Vries–Thornley Respiration Paradigms: 30 Years Later. Ann Bot. 2000;86:1–20. 10.1006/anbo.2000.1175

3. Mancuso S, Boselli M. Characterisation of the oxygen fluxes in the division, elongation and mature zones of Vitis roots: influence of oxygen availability. Planta. 2002;214:767–74. 10.1007/s004250100670

4. Arru L, Fornaciari S, Mancuso S. New Insights into the Metabolic and Molecular Mechanism of Plant Response to Anaerobiosis. Int Rev Cell Mol Biol. 2014;311:231–64. 10.1016/b978-0-12-800179-0.00005-2

5. Drew MC. OXYGEN DEFICIENCY AND ROOT METABOLISM: Injury and Acclimation Under Hypoxia and Anoxia. Annu Rev Plant Physiol Plant Mol Biol. 1997;48:223–50. 10.1146/annurev.arplant.48.1.223

6. Hebelstrup KH, Møller IM. Reactive Oxygen and Nitrogen Species Signaling and Communication in Plants. Signal Commun Plants. 2014;63–77. 10.1007/978-3-319-10079-1_4

7. Armstrong W. Aeration in Higher Plants. Adv Bot Res. 1980;7:225–332. 10.1016/s0065-2296(08)60089-0

8. King FH. Contributions to Our Knowledge of the Aeration of Soils. Science. 1905;22:495–9. 10.1126/science.22.564.495

9. Gliński J, Stępniewski W. Soil Aeration and Its Role for Plants. 2018. 10.1201/9781351076685

10. Holtman WL, Oppedijk BJ, Vennik M, Duijn B van. Low-Oxygen Stress in Plants, Oxygen Sensing and Adaptive Responses to Hypoxia. Plant Cell Monogr. 2013;371–80. 10.1007/978-3-7091-1254-0_19

11. Foley JA, Ramankutty N, Brauman KA, Cassidy ES, Gerber JS, Johnston M, et al. Solutions for a cultivated planet. Nature. 2011;478:337–42. 10.1038/nature10452

12. AlShrouf A. Hydroponics, Aeroponic and Aquaponic as Compared with Conventional Farming. American Scientific Research Journal for Engineering, Technology, and Sciences. 2017;27:247–55.

13. Barrett GE, Alexander PD, Robinson JS, Bragg NC. Achieving environmentally sustainable growing media for soilless plant cultivation systems – A review. Sci Hortic. 2016;212:220–34. 10.1016/j.scienta.2016.09.030

14. Gruda NS. Increasing Sustainability of Growing Media Constituents and Stand-Alone Substrates in Soilless Culture Systems. Agronomy. 2019;9:298. 10.3390/agronomy9060298

15. Ma L, Shi Y, Siemianowski O, Yuan B, Egner TK, Mirnezami SV, et al. Hydrogel-based transparent soils for root phenotyping in vivo. Proc Natl Acad Sci. 2019;116:11063–8. 10.1073/pnas.1820334116

16. Downie H, Holden N, Otten W, Spiers AJ, Valentine TA, Dupuy LX. Transparent Soil for Imaging the Rhizosphere. PLoS ONE. 2012;7:e44276. 10.1371/journal.pone.0044276

17. Cejas CM, Hough LA, Beaufret R, Castaing J-C, Frétigny C, Dreyfus R. Preferential Root Tropisms in 2D Wet Granular Media with Structural Inhomogeneities. Sci Rep. 2019;9:14195. 10.1038/s41598-019-50653-8

18. Benjamin AD, Abbasi R, Owens M, Olsen RJ, Walsh DJ, LeFevre TB, et al. Light-based 3D printing of hydrogels with high-resolution channels. Biomed Phys Eng Express. 2019;5:025035. 10.1088/2057-1976/aad667

19. Chen Z, Zhao D, Liu B, Nian G, Li X, Yin J, et al. 3D Printing of Multifunctional Hydrogels. Adv Funct Mater. 2019;29. 10.1002/adfm.201900971

20. Kalossaka LM, Mohammed AA, Sena G, Barter L, Myant C. 3D printing nanocomposite hydrogels with lattice vascular networks using stereolithography. J Mater Res. 2021;36:4249–61. 10.1557/s43578-021-00411-2

21. Li J, Xie J, Wu Q, Wu G, Men Y. 3D-printed hydrogel substrates with tailored pore architectures enhance root development and elicit species-specific growth responses. Chem Eng J. 2025;512:162425. 10.1016/j.cej.2025.162425

22. Ferro ND, Morari F. From Real Soils to 3D-Printed Soils: Reproduction of Complex Pore Network at the Real Size in a Silty-Loam Soil. Soil Sci Soc Am J. 2015;79:1008–17. 10.2136/sssaj2015.03.0097

23. Bacher M, Schwen A, Koestel J. Three-Dimensional Printing of Macropore Networks of an Undisturbed Soil Sample. Vadose Zone J. 2015;14:1–10. 10.2136/vzj2014.08.0111

24. Kalossaka LM, Sena G, Barter LMC, Myant C. Review: 3D printing hydrogels for the fabrication of soilless cultivation substrates. Appl Mater Today. 2021;24:101088. 10.1016/j.apmt.2021.101088

25. Ma L, Chai C, Wu W, Qi P, Liu X, Hao J. Hydrogels as the plant culture substrates: A review. Carbohydr Polym. 2023;305:120544. 10.1016/j.carbpol.2023.120544

26. Yoo D-J. Advanced porous scaffold design using multi-void triply periodic minimal surface models with high surface area to volume ratios. Int J Precis Eng Manuf. 2014;15:1657–66. 10.1007/s12541-014-0516-5

27. Han L, Che S. An Overview of Materials with Triply Periodic Minimal Surfaces and Related Geometry: From Biological Structures to Self-Assembled Systems. Adv Mater. 2018;30:e1705708. 10.1002/adma.201705708

28. Feng J, Fu J, Yao X, He Y. Triply periodic minimal surface (TPMS) porous structures: from multi-scale design, precise additive manufacturing to multidisciplinary applications. Int J Extreme Manuf. 2022;4:022001. 10.1088/2631-7990/ac5be6

29. Salvalaio M, Oliver N, Tiknaz D, Schwarze M, Kral N, Kim S-J, et al. Root electrotropism in Arabidopsis does not depend on auxin distribution but requires cytokinin biosynthesis. Plant Physiol. 2022;188:1604–16. 10.1093/plphys/kiab587

30. Soffer H, Burger DW. Effects of Dissolved Oxygen Concentrations in Aero-hydroponics on the Formation and Growth of Adventitious Roots. J Am Soc Hortic Sci. 1988;113:218–21. 10.21273/jashs.113.2.218

31. Engelmann K, Purugganan M. The Molecular Evolutionary Ecology of Plant Development: Flowering Time in Arabidopsis thaliana. Adv Bot Res. 2006;44:507–26. 10.1016/s0065-2296(06)44013-1

